# Yield Prediction Through Integration of Genetic, Environment, and Management Data Through Deep Learning

**DOI:** 10.1101/2022.07.29.502051

**Authors:** Daniel R. Kick, Jason G. Wallace, James C. Schnable, Judith M. Kolkman, Barış Alaca, Timothy M. Beissinger, David Ertl, Sherry Flint-Garcia, Joseph L. Gage, Candice N. Hirsch, Joseph E. Knoll, Natalia de Leon, Dayane C. Lima, Danilo Moreta, Maninder P. Singh, Teclemariam Weldekidan, Jacob D. Washburn

## Abstract

Accurate prediction of the phenotypic outcomes produced by different combinations of genotypes, environments, and management interventions remains a key goal in biology with direct applications to agriculture, research, and conservation. The past decades have seen an expansion of new methods applied towards this goal. Here we predict maize yield using deep neural networks, compare the efficacy of two model development methods, and contextualize model performance using linear models, which are the conventional method for this task, and machine learning models We examine the usefulness of incorporating interactions between disparate data types. We find a deep learning model with interactions has the best average performance. Optimizing submodules for each datatype improved model performance relative to optimizing the whole model for all data types at once. Examining the effect of interactions in the best performing model revealed that including interactions altered the model’s sensitivity to weather and management features, including a reduction of the importance scores for timepoints expected to have limited physiological basis for influencing yield – those at the extreme end of the season, nearly 200 days post planting. Based on these results, deep learning provides a promising avenue for phenotypic prediction of complex traits in complex environments and a potential mechanism to better understand the influence of environmental and genetic factors.

## Introduction

Prediction of an organism’s phenotype is a key challenge for biology, especially when integrating the effects of genetics, environmental factors, and human intervention. For many traits, prediction is complicated by interactions between these factors. For example, within a large multi-site, multi-genotype maize (*Zea mays*) study, more variation in grain yield is explained by interactions between genetic and environmental factors than by genetic main effects (Rogers *et al*. 2021). Including interaction effects between environmental and genomic data can improve predictive accuracy in novel environments or for new cultivars (Li *et al*. 2021; Jarquin *et al*. 2021).

Within agriculture, diverse methods have been applied to the task of predicting phenotype ranging from classical statistics (Jarquin *et al*. 2021; Rogers *et al*. 2021; Rogers and Holland 2021), machine learning (Westhues *et al*. 2021), physiological crop growth models (Technow *et al*. 2015), to combinations of these and other methods (Messina *et al*. 2018; Shahhosseini *et al*. 2021). Each model contains limitations such as lacking the capacity to model complex non-linear responses (linear models) or interactions between factors, interpretability within a biological framework (machine learning models), or dependence on costly, low throughput data for calibration (crop growth models). Often simplifying assumptions are introduced into the model (e.g. linearity), into the data (dimensionality reduction, feature engineering), or into the experimental design (e.g. considering exclusively genetic, environmental, or managerial effects to the exclusion of all others). While this approach creates more manageable statistical models and enables a sufficiently powered study to be achieved with fewer resources, it limits the capacity of a model to generalize to new genotypes, environments, or management schemes. Furthermore, which factors are treated as “nuisance” variables varies between communities within agriculture: geneticists often restrict management regimes, while agronomists usually consider only a few cultivars. These approaches make it difficult to investigate the interactions between genetic, environmental, and management factors.

To predict an organism’s phenotype across genotypes, environments, and management strategies simultaneously requires a dataset containing many combinations of these features.

Collecting such a dataset requires a large multi-site, multi-condition, experiment featuring diverse genetic backgrounds. The Genomes to Fields Initiative (McFarland *et al*. 2020) seeks to accomplish this aim. To date it has collected measurements of grain yield and other phenotypic traits (plant height, days to silking, stalk lodging, and kernel row number, etc.) from about 180,000 plots planted at more than 160 environments. Environments are characterized using a WatchDog 2700 Weather station (Spectrum Technologies, Inc.) which collects continuous weather data thought the season and collaborator submitted soil samples. Across the initiative, over 2,500 maize hybrids have been tested, with Genotyping by Sequencing performed on inbred parental lines used. Beyond the data collection, a means of effectively incorporating diverse data types (genomics, management, soil measurements, weather, etc.) is needed, particularly one that avoids simplifying assumptions where possible.

One method with the potential to accomplish this is that of deep neural networks (DNNs) which have the capacity to approximate any function, provided they are sufficiently complex and have sufficient examples to learn from. This capability is present regardless of whether they are composed of dense fully connected (Hornik *et al*. 1989) or convolutional layers (Zhou 2020).

Additionally, DNNs “learn” directly from the data provided which enables reduced feature engineering and dimensionality reduction. The methodology is also flexible with respect to data type, allowing combination of variables that are static over a growing season (e.g. genotype) and those that are dynamic (e.g. temperature) in a single model (Washburn *et al*. 2021). While neural networks have been applied to the problem of predicting yield since at least 2001 (J. Liu *et al*. 2001) this field is rapidly developing, with advances in theory, software, and hardware enabling deeper and more accurate networks. Several recent studies have applied these methods with a relatively large dataset either with (Washburn *et al*. 2021) or without (Khaki *et al*. 2020) a genetic component into the model, with little feature engineering performed. Both relied instead on DNN’s capacity to learn useful data transformations from the data directly.

Despite their promise, DNNs are not a panacea for prediction. DNNs are prone to overfitting to training data resulting in poor performance. Even when performing well, the complexity of these models can obscure what aspects of the data the model is using. Advances in deep learning have produced methods which reduce these limitations. For example, the use of convolutional layers minimizes the potential of overfitting because they perform well with fewer parameters relative to fully connected layers. Where fully connected layers are used, overfitting can be reduced by randomly removing neurons from a layer with a certain “dropout” percentage. While the inner workings of DNNs remain far less interpretable than simpler models (e.g., Genomic BLUP or physiological models), methods have been developed to aid in interpretation through identifying the importance of different features in the data which can be applied. These methods include salience (Simonyan *et al*. 2014), guided backpropagation (Khaki *et al*. 2020), and permutation based metrics (Shahhosseini *et al*. 2021) among others (Samek *et al*. 2017). Here we use salience to illuminate the operation of the DNNs generated in this study.

Here we, leverage DNNs’ capacity to determine feature importance from the data which permits us to remain agnostic as to which features, or combinations of features are most relevant. Furthermore, since DNNs are robust to lower--quality data and benefit from an abundance of data, we employ a strategy of minimal feature transformation and curation and maximal inclusion of observations. Using a minimally transformed dataset we begin the search space considered in (Washburn *et al*. 2021), expand the space under consideration, and detail a sequence of reproducible steps and objective heuristics which produced the models under consideration. DNNs require an abundance of data for training. We begin by detailing a workflow that incorporates a wider number of years from the Genomes to Fields Initiative in than previous studies (Rogers *et al*. 2021; Washburn *et al*. 2021; Rogers and Holland 2021), while also limiting the effect of errant and absent measurements. Improving on past studies, we propose a new approach to model optimization whereby the model is broken into sub-modules for each data type and interactions between them, then each submodule is consecutively optimized, using a bayesian optimization procedure to find a suitable structure based on the data itself. As far as we are aware, previous studies using deep learning for phenotypic prediction have instead employed simultaneous optimization of all model components (Washburn *et al*. 2021) or informal inductive tinkering. We compared models developed through consecutive and simultaneous optimization and tested them against a variety of classic machine learning and statistical methods to determine which performed best. To fairly assess model performance we detail a strategy of constructing testing, training, and validation sets stratified by season and location that is broadly useful to assessing model performance, while avoiding overfitting the model to any location

## Materials and Methods

### Data Preparation

We used data from the Genomes to Fields (G2F) initiative for years 2014-2019 (McFarland *et al*. 2020), focusing on the sites within the continental United States. Each year’s data are publicly available (https://www.genomes2fields.org/resources/), including weather and soil data for field sites, genomic data, management schedules (e.g., application of fertilizer, herbicides, irrigation) and yield (in addition to other phenotypic variables). We augmented this through additional genomic and weather data. Weather data retrieved from Daymet (Thornton *et al*. 2020) was used in quality control as discussed below and to infer data for locations which lacked a functional weather station for some or all of the season. Daymet data was retrieved through wget (Techtonik 2015).

These data are provided with some variability in format. Custom scripts were used to aggregate and standardize terminology across years. Rather than itemizing each operation, we restrict ourselves to those which are likely to be of interest to those working with similar data sets. The scripts used are available through Bitbucket (https://bitbucket.org/washjake/maizemodel and https://bitbucket.org/daniel_kick/maizemodel/ ).

Scripts were written in Python (Van Rossum and Drake 2009 p. 3) and rely on scientific and common general libraries (Seabold and Perktold 2010; Pedregosa *et al*. 2011; *fuzzywuzzy* 2017; Virtanen *et al*. 2020; team 2020; Harris *et al*. 2020; Da Costa-Luis *et al*. 2022) along with plotting libraries for exploratory visualizations (Hunter 2007 p. 200; Inc 2015; Waskom 2021; Kibirige *et al*. 2021). We used Anaconda (“Anaconda Software Distribution” 2021) to manage the virtual environment.

We reduced the dimensionality of the genomic data with principal components analysis (PCA) before use. Provided genomes were loaded into TASSEL version 5.2.74 (Bradbury *et al*. 2007) and filtered. Filter parameters were intended to be relatively unrestrictive, while still reducing the data enough for PCA to complete with the memory available. We arrived at these filtering parameters through inductive tinkering (i.e., iteratively increasing filtering until the dataset was sufficiently small for the process to not exhaust available memory). First, we restricted the loci to those having a heterozygosity of at least 0.001. Next, we converted the data to a numerical genotype, and imputed missing values are imputed with the mean. The quality of observations varied through the dataset. We discarded samples if they had missing values for 90% or more of loci. Once the data was reduced, the genomes were PCA transformed. We find that 31% of the variance is explainable by the first 8 principal components (PCs), 50% is explainable by the first 50 PCs, and >99% of the variance is explainable by 1725 PCs.

Each hybrid’s coordinates in PC space were estimated as the average between its parent’s coordinates. This was done rather than creating simulated hybrids due to hardware and software constraints. If simulated hybrids are generated in TASSEL the number of observations input into PCA is substantially increased (i.e., all observed combinations of genomes are transformed, not all observed genomes). Completion of PCA would thus requires stricter filtering for the analysis to be completed. By estimating hybrid values after PCA transformation we retain a greater number of loci per observation influencing the PCs.

Environmental data required preprocessing as well. The soil dataset contains many missing values, having an average completion rate of 47% across all site-by-year combinations. For each variable in the soil dataset, missing values were first linearly interpolated across years with respect to location. Locations with no observations for any years were imputed using k-nearest neighbors based on the nearest 5 neighbors for remaining missing values. Within the reported weather data, we observed evidence of equipment malfunction and imputed or adjusted values using linear models.

First, we removed outliers and extreme observations by limiting the data to those days where the difference between the minimum daily value estimated by Daymet and the measured value were in the central 60% of the distribution. This removes values from the G2F reported weather data (based on more inexpensive and error prone equipment) that are inconsistent with the Daymet data. Next, linear models were generated using the 4 Daymet estimated variables with the strongest correlations with the target variable across the dataset. All combinations of additive models containing 0-4 of these predictors plus an optional site-by-year term were fit and ranked using Akaike information criterion corrected for small sample size (AICc). The best performing model for each metric was used to impute missing weather data. In the case of temperature measurements, which exhibited the most apparent evidence of instrumental error, observed values were also replaced if the difference between the expected and observed value was greater than 1.5x the interquartile range of those differences.

The representation of management data was refined. Fertilizer applications were decomposed into the quantity of nitrogen, phosphorus, and potassium applied. Where fertilizer applications were lacking an application date, we estimated the time difference relative to the planting date with K-NN imputation (k = 5) to cluster based on application quantity. To define the time window under consideration, we used the earliest within-season fertilizer application and the day of the latest harvest to bound the weather and management data. This resulted in selecting 75 days prior to planting, 1 planting day, and the 204 subsequent days (210 total days).

Weather and management time-series data were clustered to reduce their dimensionality for use in machine learning and linear models. For each variable we used time series k-means with dynamic time warping implemented through the tslearn library (Tavenard *et al*. 2020). K was optimized by calculating the silhouette score for k between 2 and 40 then selecting the k one less than the lowest k in which the silhouette score decreased. Where needed clusters were represented categorically through one hot encoding.

### Defining Training, Validation, and Test sets

We generated train/test splits randomly, with the constraint that any location-year combination could appear in only the testing or training set. Nearby experimental sites were grouped for the purpose of generating training and testing sets. The method of generating splits was: (1) one site group was selected at random and added to the testing set. (2) Each site-group-by-year combination was down sampled so that it had no more observations than were found in the smallest site-group-by-year combination in the testing set. This prevented overrepresentation of any one group in the training data. (3) This process was repeated until the testing set accounted for 10% to 15% of the total observations *and* at least 40,000 observations remained across the training and test set. If these conditions were not met the process was repeated with a different seed value.

During deep neural network hyperparameter selection, we limited overfitting to a single validation set by using Monte Carlo cross-validation, stratified by site-group-by-year. Folds were created while controlling for year by location groups by drawing the same number of site-group-by-year groups as were present in the test set (3 for the test/train split used here) but without further down sampling. The selected groups constituted the validation set. This was repeated 10 times and the membership of the folds preserved throughout training or hyperparameter search for a single model. To enable reproducibility, random number generator seeds were used in train/validate split searching and Monte Carlo cross-validation fold generation.

Prior to hyperparameter selection and training the input data was centered and scaled based on the mean (∼147.397 bushels per acre) and standard deviation (∼48.169 bushels per acre) of the yield in the training data, i.e., 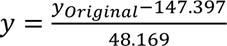 Transforming the data prior to determining cross validation folds potentially introduces an information leak within the hyperparameter selection process (i.e., the data used in evaluation, here across cross validation folds, by influencing the mean and standard deviation used in centering and scaling the data).

However, this does not create an issue for final evaluation of the model’s performance because the test set data were not used in these calculations.

### Model Preparation

#### Overview

We sought to model genotype by environment by management interaction effects (GEM effects) in maize yield and to determine utility of doing so. To this end we optimized DNNs to predict yield with a single data modality (i.e., only genomic data, soil characteristics, or time series data each by itself). We use one dimensional convolutional layers to capture the time dependent features of weather data, which have previously been used in yield prediction for this task (Khaki *et al*. 2020; Washburn *et al*. 2021). We used dense, fully connected layers for the other submodules of the DNN.

We pursued two strategies for tuning and training GEM models: Consecutive Optimization (CO) and Simultaneous Optimization (SO). CO tunes the hyperparameters of networks predicting yield from a single data modality (genomic data, soil data, or weather and management time series data). Next, the prediction neurons are discarded and the output of the penultimate layer of each single modality network enters a set of layers to permit interactions between data modalities. Hyperparameters for the interaction layers are then tuned. The SO strategy by contrast allows for all hyperparameters to be selected concurrently, both those which affect the processing of a single data modality and those influencing interactions between modalities.

#### Hyperparameter Search and Training

We selected model architecture through a hyperparameter search using the ‘BayesianOptimization’ tuner provided within the ‘keras-tuner’ package (O’Malley *et al*. 2019). Models were written in Keras (Chollet and others 2015) with Tensorflow as a backend (Martín Abadi *et al*. 2015) and run in a Singularity container (Kurtzer *et al*. 2017; SingularityCE Developers 2021). The subnetworks processing exclusively genomic and exclusively soil data, along with the interaction subnetwork, are constructed exclusively of dense (i.e., fully connected) layers, each subject to batch normalization and dropout. The weather/management processing subnetwork is composed of two one-dimensional convolution layers followed by batch normalization and a pooling layer. The output of this subnetwork is flattened before entering the interaction subnetwork. Hyperparameter ranges explored for each network are listed in Table 1. To avoid overfitting to the validation data we used a custom subclassed version of the tuner to randomly select one of the previously defined validation folds. This is done rather than using mean loss over all folds to avoid increasing the computational cost 10- fold while still preventing overfitting to a single validation set.

**Table 1.**
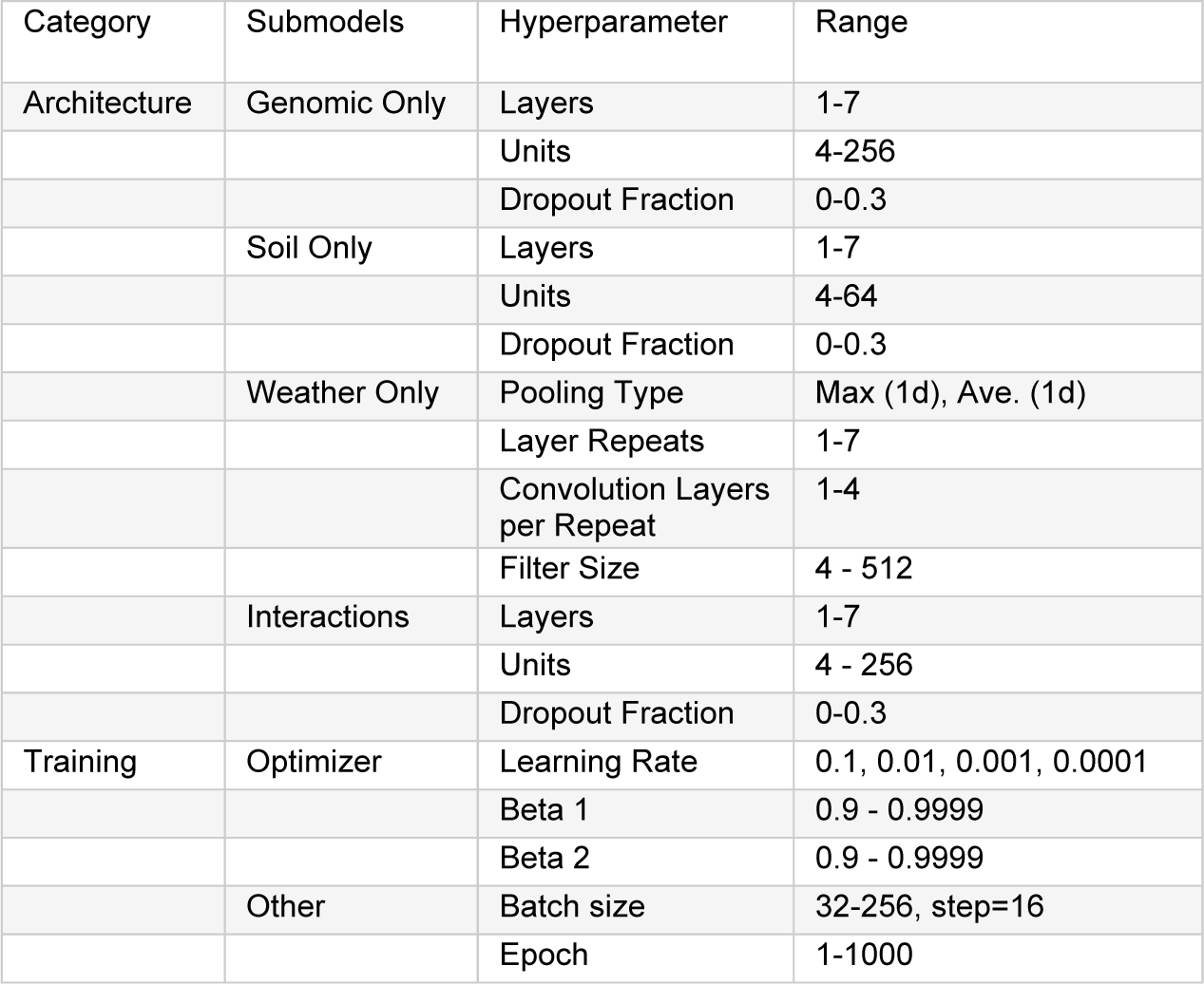
Hyperparameter Ranges: Deep Learning

For all DNNs, a maximum of 40 hyperparameter sets were explored. In cases where no convolution layers were being varied (CO models with only genomic, or only soil data, and the interaction layers trained for the same strategy) hyperparameters were trained for a maximum of 1000 epochs with an early stopping patience of 7 epochs or more. For cases in which convolution layers were varied (Sequential Optimization model with only weather and management data, Concurrent Optimization model) hyperparameters were trained for a maximum of 500 epochs with an early stopping patience of 5 epochs. This difference is due to practical rather than theoretical reasons as the convolutional networks required notably more time per epoch to fit. Regardless of network type, if the hyperparameters optimization had not concluded by 290 hours after the script began, the process was terminated and the hyperparameter sets completed by that point were considered.

To ensure the selected hyperparameter set performs well across validation sets, the top 4 hyperparameter sets for each model were trained for 1000 epochs and evaluated on all 10 defined testing/validation set splits. Next, the validation losses over the duration of training were used to calculate the mean and standard deviation for each epoch. Then the training duration was split into 10 bins and the average of the sum of validation loss mean and standard deviation was calculated, i.e. 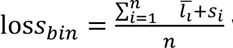 where *i* is epoch relative to the beginning of the bin, 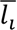 is the mean validation loss across cross validation folds at the i^th^ epoch and *s*_*i*_ is the standard deviation of the same. The hyperparameter set with the lowest value for the most bins was used going forward.

For the best hyperparameter set, we selected a training duration from the validation losses. For each fold we calculated a rolling mean of validation loss with a window size of 20 epochs. Next, for each epoch we calculated the sum of the mean and standard deviation of the rolling mean and the total rolling validation loss. Then we found the epochs which minimized these two values (subtracting 10 from the epoch number to account for the window size). The disagreement between the epochs which minimized these values ranged from 2 epochs in the case of the CO Genomic model and CO interaction model up to 404 epochs for the CO weather and management model. We decided to use total rolling validation loss to decide on the epoch number for each model. This metric resulted in more training epochs for all models except the SO model. Incorporating a more sophisticated method for selecting training duration is a possible improvement for future studies. With the selected hyperparameters and training duration we fit each model 10 times to account for random initialization and saved each replicate and its training history.

### Benchmarking Models Overview

To contextualize the performance of the generated deep neural networks we use the same training data to fit linear and classic machine learning models. These models often require fewer resources and time to train than deep neural networks. For linear models we consider a small collection of models that varied with respect to the independent variables present, whether interactions are included, and whether effects are fixed or random. For supervised machine learning models, we selected and optimized four methods: k-nearest neighbor (KNN), radius neighbor regression (RNR), random forest (RF), and support vector regression with a linear kernel (SVR).

#### Linear Models

To aid in evaluating the efficacy of the models produced, we constructed linear models varying in the scope of included data and model complexity. The simplest model was an intercept model, i.e., every predicted yield equals the mean yield in the training set. We considered three models using the genomic data alone: fixed effects for PCs 1-8 (31% variance explained), fixed effects for PCs 1-50 (50% variance explained), and random effects for the first 8 PCs. For soil data we considered two models, one with all factors as fixed effects and one with all factors as random effects. From weather and management data we produced three models, using all factors as fixed effects, using clusterings of the top five most salient features (i.e., Water total, solar radiation mean, maximum temperature, mean wind direction, vapor pressure) identified in the weather and management data (averaging over time points) as fixed effects, and as random effects. Most salient features were taken from the deep neural network with the lowest average test set RMSE (CO Interaction). All weather and management data were represented as categorical clusters as described in “Data Preparation”.

We evaluated five models using a combination of data sources. In three fixed effect models we incorporated: (1) PCs 1-8 and the five most salient weather features’ clusterings, (2) the same plus all soil features, and (3): the effects in “1” plus interactions between each PC and weather factor cluster. The two random-effects models we fit using PCs 1-8 and the selected weather clusters excluding or allowing interactions. We fit Fixed effect models with the linear model function in R (R Core Team 2021) and random effect models with lme4 (Bates *et al*. 2015). This analysis was aided by common data wrangling and convenience libraries (Wickham *et al*. 2019; Bache and Wickham 2020; Müller 2020; Izrailev 2021) and feather file read/write capabilities through arrow (Richardson *et al*. 2021).

#### Classical Machine Learning Models

Additional machine learning models were implemented through scikit-learn (Pedregosa *et al*. 2011; Buitinck *et al*. 2013) and hyperparameters for each were optimized through the hyperopt library (Bergstra *et al*. 2013) run within a Docker container. In a workflow similar to that of the deep neural network models, we generated models for each data modality indpendently, and with all data available. Time series data was represented as clusters as described in “Data Preparation”. For each model we allowed the following hyperparameters to vary as described: (1) K Nearest Neighbors (KNN): neighbors = 1-250, weights = ’uniform’ or ’distance’; (2) Radius Neighbors Regressor (RNR): radius = 0.01-2000, weights = ’uniform’ or ’distance’; (3) Random Forest (RF), maximum depth = 2-200, Minimum samples per leaf = 0-0.5; and (4) Support vector machine with a linear kernel (SVR): Loss = ’epsilon_insensitive’ or ’squared_epsilon_insensitive’, C =1-5 (log uniformly distributed).

Cross validation folds matching those as described previously and average loss across all folds was measured. We tested a minimum of 115 combinations for each model and selected the best performing hyperparameters for each input dataset, reported in Table 5. Following selection, we trained each model and produced predictions on the testing and training data. This was repeated 10 times to account for randomness in model fitting.

### Model evaluation

For every model described above we calculate predicted yields for the test set and calculate root mean squared error 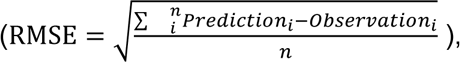 , normalized RMSE percent 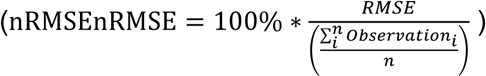 and R^2^ using SciPy (Virtanen *et al*. 2020).

Unless stated otherwise in the text RMSE and nRMSE will refer to the average value across replicates. Two observations were not predictable using the fit radius neighbors regressor and were predicted as the training set mean. For the best performing DNN we calculated and visualized the salience of features for each data modality. To examine the influence of allowing interactions we contrast these saliences with the saliences of SO single modality DNNs.

Saliences were calculated by Tf-keras-vis (Kubota 2021). Visualizations were created with the use of rjson (Couture-Beil 2018), patchwork (Pedersen 2020), and ggplot2 (Wickham *et al*. 2019).

### Data Availability Statement

Data for maize phenotypes, genotypes, field site soil properties and on location weather recordings from the Genomes to Fields Initiative data (McFarland *et al*. 2020) is publicly available through the CyVerse Discovery Environment. We used data from 2014 to 2019 which correspond to the following DOIs: 2014 – 2017 (0.25739/frmv-wj25), 2018 (10.25739/anqq- sg86), and 2019 (10.25739/t651-yy97). Additional genomic data was provided by Natalia de Leon, Dayane Lima, and Cinta Romay worked with Joseph Gage through personal communication. These data will be available through CyVerse. Following public release, a version of these data containing all genomic data used in this study will be available through Zenodo (10.5281/zenodo.6916775). At time of writing, this repository contains the eigenvectors resulting from the principal components analysis are provided to enable transformation of provided genomes. Additional weather measurements were retrieved from Daymet (Thornton *et al*. 2020). Custom python scripts for downloading, aggregating and processing these data are available on bitbucket (https://bitbucket.org/washjake/maizemodel and https://bitbucket.org/daniel_kick/maizemodel/ ) in the notebooks directory (files with the prefix 0.0 to 0.5).

## Results

### Deep Neural Networks can--but do not necessarily--outperform competing model types

When all data sources are incorporated, the CO DNN achieves the best (i.e., lowest) average RMSE (when not otherwise specified, values refer to the average across replicates), followed by a fixed effect model with interaction effects (nRMSE 14.6% vs 14.9%, RMSE 0.948 vs 0.959) (Figure 2, Table 6). Despite this, with 2/10 replicates of the CO DNN model underperform this fixed effect model. Variability in DNN performance across replicates can be caused by the random initialization of the weights at the start of training.

Following CO DNN and a linear model with interaction effects, an exclusively additive linear model incorporating all data modalities ranked third (nRMSE 15.0%, RMSE 0.980) and one excluding soil data ranked fourth (nRMSE 15.1%, RMSE 0.981). Random effects models with and without soil data (nRMSE 15.2%, 15.3%, RMSE 0.991, 0.994) and SO DNN (nRMSE 15.7%, RMSE 1.024) followed. Of the machine learning models only support vector regression with a linear kernel (SVR) and K Nearest Neighbor (KNN) outperformed a simple intercept model. We find similar results for R^2^ (Supplemental Figure 2, Table 6).

A DNN is not the best performing model when data is restricted to a single modality.

When restricted to genomic data, KNN, followed by linear models with fixed or random effects are the only ones which outperform the intercept model (nRMSE 16.5%, 16.6%, 16.7%, 16.7%, RMSE 1.078, 1.084, 1.085, and 1.088, KNN, linear fixed effects, linear random effects, intercept model). SVR performed particularly poorly on this data (nRMSE 18.7%, RMSE 1.212) -- nRMSE 2% or RMSE of 0.131 above the intercept model. Incorporating only soil data SVR performed best (nRMSE 16.3%, RMSE 1.059) with all other models being within nRMSE 0.261% or 0.017 RMSE of the intercept model. Most models performed better when instead trained on weather/ management data with the exception of the random forest (RF) which had an nRMSE 5.729%, RMSE of 0.373 *above* the intercept model. SVR (nRMSE 15.1%, RMSE 0.985) and a fixed effects model (nRMSE 15.2%, RMSE 0.993) performed remarkably well. CO DNN is capable of outperforming these methods, but not uniformly. 2/10 replicates underperformed the intercept model resulting in a nRMSE 15.6%, RMSE of 1.018 while the median values were nRMSE 15.2% and RMSE is 0.992.

### Consecutive Optimization resulted in a larger, more accurate final network

Two hyperparameter selection strategies were employed, Consecutive Optimization (CO) and Simultaneous Optimization (SO), have the same range of possible networks (hyperparameter ranges are listed in Table 1), the same data driving network selection and both use bayesian optimization. Despite this, the strategy applied resulted in notably different final architectures. A visual summary of the relative differences between network hyperparameters is shown in Figure 1, with the hyperparameter values listed in Tables 2 and 3. Supplementary Figure 1 provides a visual overview of the network architecture. We consider the effect of CO vs SO on each of the four subnetworks (processing exclusively genomic, soil, or weather/management factors or interactions between data modalities), listed in decreasing order of approximate similarity.

**Table 2.** Selected Deep Learning Hyperparameters: Architecture

| Submodel or Network | Hyperparameter | Specific Layer | Consecutive Optimization | Sequential Optimization |
| --- | --- | --- | --- | --- |
| Genomic Only | Units | 1 | 83 | 196 |
|  |  | 2 | 133 | 47 |
|  | Dropout Fraction | 1 | 0.163923177 | 0.15214 |
|  |  | 2 | 0.230663142 | 0.06061 |
| Soil Only | Units | 1 | 38 | 19 |
|  |  | 2 | 13 | 27 |
|  |  | 3 | 45 |  |
|  |  | 4 | 29 |  |
|  |  | 5 | 4 |  |
|  |  | 6 | 4 |  |
|  |  | 7 | 4 |  |
|  | Dropout | 1 | 0.148724301 | 0.21342 |
|  |  | 2 | 0.276340999 | 0.18589 |
|  |  | 3 | 0.005434164 |  |
|  |  | 4 | 0.173380695 |  |
|  |  | 5 | 0 |  |
|  |  | 6 | 0 |  |
|  |  | 7 | 0 |  |
| Weather + Management Only | Pooling Type | N/A | Max | Max |
|  | Convolution Layers per Repeat | N/A | 2 | 2 |
|  | Filter Size | 1 | 433 | 370 |
|  |  | 2 | 436 | 303 |
|  |  | 3 | 52 |  |
|  |  | 4 | 163 |  |
|  |  | 5 | 400 |  |
|  |  | 6 | 294 |  |
| Interaction | Units | 1 | 152 | 10 |
|  |  | 2 | 207 | 25 |
|  |  | 3 | 206 | 126 |
|  |  | 4 | 188 | 204 |
|  |  | 5 | 44 | 45 |
|  |  | 6 |  | 134 |
|  | Dropout | 1 | 0.18658661 | 0.10201 |
|  |  | 2 | 0.289893588 | 0.14809 |
|  |  | 3 | 0.004841293 | 0.01536 |
|  |  | 4 | 0.198121953 | 0.15658 |
|  |  | 5 | 0.243027717 | 0.2428 |
|  |  | 6 |  | 0.19048 |

**Table 3.** Selected Deep Learning Hyperparameters: Training

|  | Optimizer |  | Other |  |  |
| --- | --- | --- | --- | --- | --- |
| Network | learning_rate | beta1 | beta2 | numEpoch | batch_size |
| CO: Genomic Only | 0.0001 | 0.953368 | 0.985947 | 12 | 96 |
| CO: Soil Only | 0.01 | 0.928472 | 0.997516 | 199 | 176 |
| CO: Weather Only | 0.0001 | 0.903649 | 0.929582 | 629 | 240 |
| CO: Full Network | 0.01 | 0.98752 | 0.972311 | 364 | 112 |
| SO | 0.001 | 0.975893 | 0.994607 | 711 | 192 |

**Figure 1.**
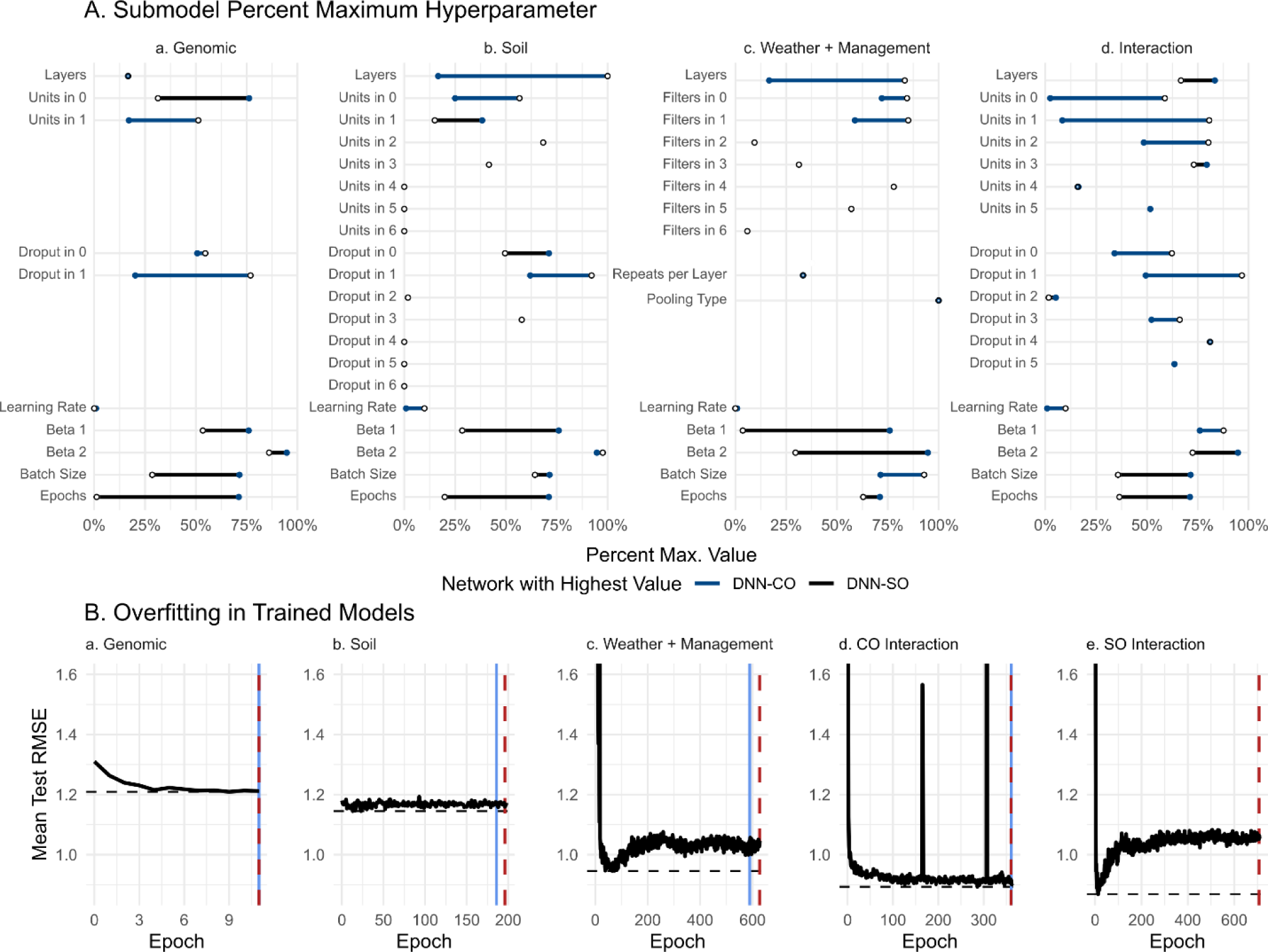
Optimization strategy results in different network architectures and degree of overfitting in full model **A.** Hyperparameters for each optimization strategy are shown as a percent of the allowed range. Data available for training is the same but the Consecutive Optimization (CO) and Simultaneous Optimization (SO) strategy result in substantially different hyperparameter values and thus network architecture. For exact values refer to Table 2 and Table 3. **B.** The average RMSE of the test set (across 10 replicates to account for random initialization of weights) is shown in black for each submodel (**a. – e.**). The horizontal dashed black line indicates the minimum error achieved throughout the training duration. The vertical lines indicate the difference in error and epochs of the minimum value and the values selected through minimizing total validation error (red dashed line), the heuristic used in this study, and the mean plus standard deviation of validation error (solid blue line), which was considered but not used. Both strategies considered failed to select the epoch resulting in the minimum loss in the test set for all submodels and resulted in apparent overfitting in the Weather and Management submodel (**c.**) and the SO model (**e.**). For additional comparisons of heuristic performance see Table 4.

**Table 4.**
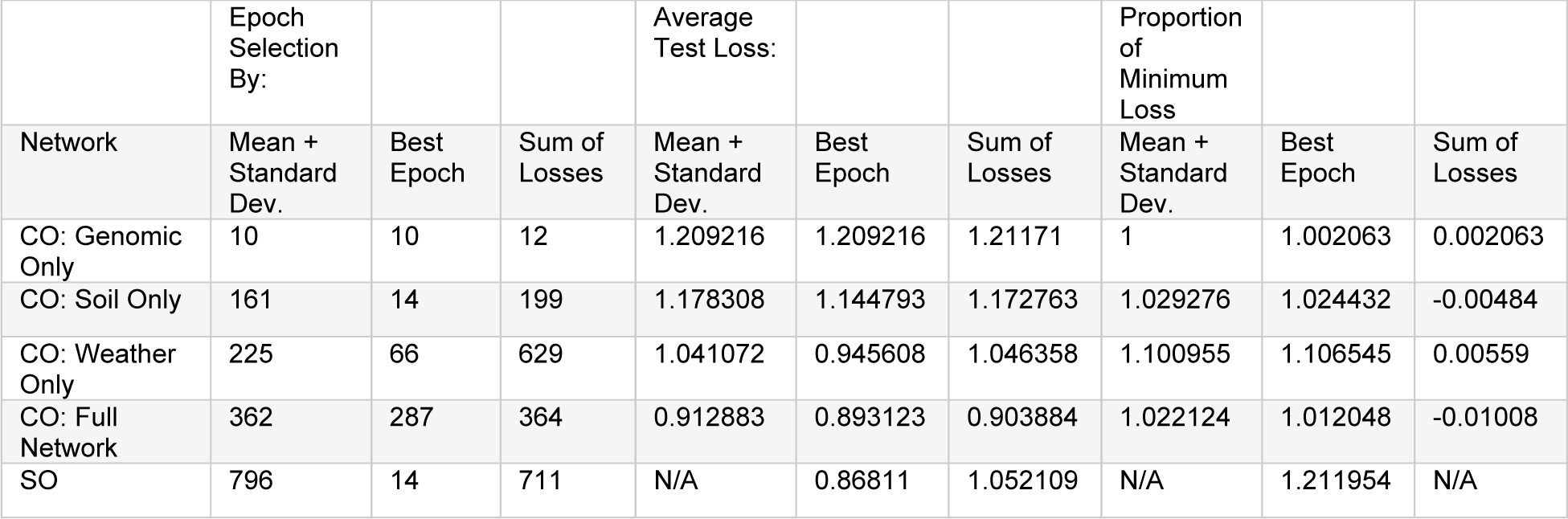
Epoch Selection Underperformance

|  | Epoch<br>Selection<br>By: |  |  | Average<br>Test Loss: |  |  | Proportion<br>of<br>Minimum<br>Loss |  |  |
| --- | --- | --- | --- | --- | --- | --- | --- | --- | --- |
| Network | Mean +<br>Standard<br>Dev. | Best<br>Epoch | Sum of<br>Losses | Mean +<br>Standard<br>Dev. | Best<br>Epoch | Sum of<br>Losses | Mean +<br>Standard<br>Dev. | Best<br>Epoch | Sum of<br>Losses |
| CO: Genomic<br>Only | 10 | 10 | 12 | 1.209216 | 1.209216 | 1.21171 | 1 | 1.002063 | 0.002063 |
| CO: Soil Only | 161 | 14 | 199 | 1.178308 | 1.144793 | 1.172763 | 1.029276 | 1.024432 | -0.00484 |
| CO: Weather<br>Only | 225 | 66 | 629 | 1.041072 | 0.945608 | 1.046358 | 1.100955 | 1.106545 | 0.00559 |
| CO: Full<br>Network | 362 | 287 | 364 | 0.912883 | 0.893123 | 0.903884 | 1.022124 | 1.012048 | -0.01008 |
| SO | 796 | 14 | 711 | N/A | 0.86811 | 1.052109 | N/A | 1.211954 | N/A |

**Table 5.** Machine Learning Hyperparameter Optimization

| Model | Hyperparameter | Range | Genomic Only | Soil Only | Weather + Management Only | Multiple |
| --- | --- | --- | --- | --- | --- | --- |
| KNN | Weight Metric | Uniform, Distance | Uniform | Distance | Uniform | Distance |
|  | k | 1 - 250 | 237 | 248 | 248 | 49 |
| RNR | Weight Metric | Uniform, Distance | Distance | Distance | Uniform | Distance |
|  | Radius | 0.01 - 2000 | 39.759518 | 3.406197 | 5.986679 | 40.375418 |
| SVR | Loss | Epsilon Insensitive, Squared Epsilon Insensitive | Epsilon Insensitive | Epsilon Insensitive | Epsilon Insensitive | Squared Epsilon Insensitive |
|  | C | 1 - 5 (log uniform) | 2.772318 | 5.613996 | 4.623351 | 2.787589 |
| RF | Max Depth | 2 - 200, q = 1 (q uniform) | 64 | 10 | 102 | 7 |
|  | Min Samples/Leaf | 1 - 200, q = 1 (q uniform) | 171 | 163 | 100 | 149 |

**Table 6.**
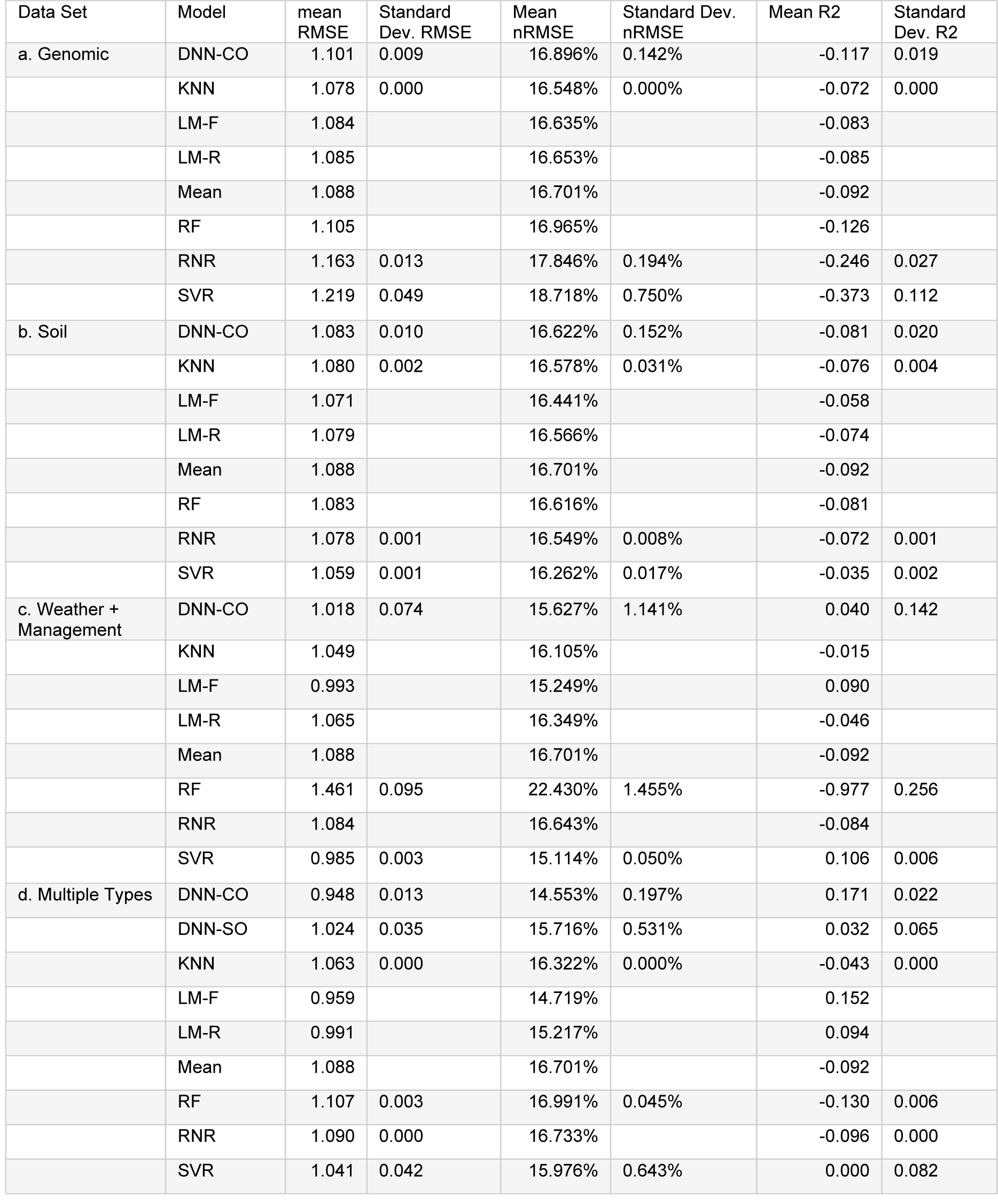
Performance Across Data Sets

The output of this subnetwork is flattened before entering the interaction subnetwork genomic subnetworks resulting from CO and SO are both two layers, but the CO model widens somewhat (layer 1 = 83 units, 16% dropout, layer 2 = 133 units 23% dropout) while the SO model begins over twice as wide and constricts more (layer 1 = 196 units, 15% dropout, layer 2 = 47 units 6% dropout). The interaction subnetworks contained a similar number of layers (CO: 5 vs SO: 6), but while CO resulted in layers with similar widths before constricting at the last layer (units = 152, 207, 206, 188, 44, dropout percentages = 19%, 29%, 0.5%, 20%, 24%), SO resulted in layers with very few units initially which are later expanded (units = 10, 25, 126, 204, 45, 134, dropout percentages = 10%, 15%, 2%, 16%, 24%, 19%). The soil subnetwork resulting from CO is notably deeper than the one from SO (7 and 2 dense layers respectively) but also narrows more by the last processing layer (2 vs 27 units). Finally, in the weather and management subnetwork CO resulted in a notably deeper network (6 pairs vs 2 pairs of convolution layers) but used a similar number of filters in the final convolution layer pairs (CO 294 vs SO 303).

The performance of these networks differs as well. The CO network was better at predicting yield in the testing set. It achieved a lower mean RMSE (CO: 0.948 vs SO: 1.024) and was more consistently accurate across replicates (standard deviation CO: 0.013 vs SO: 0.035). Similar results were seen in the normalized errors (nRMSE CO: 14.6% SO: 15.7%, standard deviation CO: 0.197%, SO: 0.531%). Similarly, average R^2^ was higher in the CO network (CO: 0.171 vs SO: 0.032) and more consistent across replicates as well (standard deviation CO: 0.022 vs SO: 0.065).

Model performance differences are due, in part, to the heuristic used to select the number of training epochs and different tendencies for these models to overfit. The heuristic used to select the number of training epochs (sum of the rolling validation loss) and alternate heuristic considered (mean plus standard deviation of the rolling validation loss) resulted in networks with comparable performance, having on average 0.001 less RMSE. With the exception of the SO DNN, this also resulted in longer training durations. These ranged from an additional 2 epochs in the cases of the CO genomic and interaction models and as many as 404 epochs in the case of the CO weather/management, as shown in Table 4.

These training durations were often considerably longer than the optimal values as seen in Figure 1B. Furthermore, the length of overtraining appears loosely proportional to the present minimum average RMSE each model achieved. The SO and CO weather models had the largest differences between optimal and used epoch numbers – differences of 697 and 563 epochs respectively and achieved 121% and 110% of the minimum possible RMSE. The CO soil model trained an excess 185 epochs but only had RMSE at 102% minimum. The two training durations closest to the optimum were the CO genomic model (2 epochs over) and the SO model (77 epochs over). These models performed at just 100.2% and 101% minimum.

The SO model overfits faster and to a greater extent than the full CO model, which does not show evidence of substantial overfitting (Figure 1B d, e). The SO model achieves a loss lower than the CO model, and the accuracy worsens rapidly with further training. The different network sizes (CO containing more layers) may account for this difference. Improved heuristics for training duration could represents an opportunity for future refinements, which these results suggest could both increase goodness of fit and reduce the computational resources needed to train these models.

### Model performance generally improves through incorporating multimodal data and interactions

Incorporating multiple data sources and allowing interactions between data types generally appears to improve accuracy. Within the tested DNNs, allowing interactions increased performance relative to single modality models. The potential exception to this is the SO DNN (nRMSE 15.7%, RMSEs 1.024) and the CO weather/management model (nRMSE 15.6%, RMSE 1.018). Despite this, the former’s distribution had lower dispersion with a standard deviation of RMSEs 0.035 relative to 0.074. Within the linear models tested, allowing interactions increased accuracy in the fixed effect model by 0.3% nRMSE or 0.023 RMSE but *decreased accuracy* in random effects models by 0.04% nRMSE or 0.003 RMSE.

In purely additive linear models, incorporating additional data modalities decreases error.

The largest difference in fixed effect models is for genomic data (improvement of 1.916% nRMSE, 0.125 RMSE), followed by soil data (improvement of 1.722% nRMSE, 0.112 RMSE), and weather/management data (improvement of 0.531% nRMSE, 0.035 RMSE). The same trend is seen in models with random effects models, albeit with less variation in improvements (improvements of 1.436% nRMSE, 0.094 RMSE genomic, 1.349% nRMSE, 0.089 RMSE soil, 1.132% nRMSE, 0.074 RMSE weather/management).

This pattern does not hold for the machine learning methods tested. For KNN, the model trained on exclusively weather data performed best (0.218% nRMSE, 0.014 RMSE better than using all data) although using all data sources did improve accuracy relative to only genomic or soil data. SVR follows a similar pattern but is more exaggerated with using exclusively weather data resulting in an improvement of 0.862% nRMSE or 0.059 RMSE relative to all data whereas all data represented an improvement relative to genomic and soil data. Random forests did not follow this trend– Genomic and soil models performed better than all data by 0.026% and 0.375% or 0.002 and 0.024 RMSE respectively, whereas the weather and management model performed 5.439% nRMSE or 0.354 RMSE worse. Finally, radius neighbor regression (RNR) performance was worst using only genomic data (1.112% nRMSE, 0.072 RMSE worse than all data) but using only soil or only weather data improves model accuracy by 0.185% and 0.090% nRMSE or 0.012 and 0.006 RMSE respectively.

### Which factors are most important to the CO DNN?

Among the genomic data we observe no major trend in salience with respect to PC (Supplementary Figure 3 A.). The two most salient PCs are PC 26 (0.423) and PC 24 (0.402) which account for 0.350% and 0.392% of the total genomic variance respectively. Given that these saliences are relative to principal components, using salience to implicate specific genes or gene loci is infeasible. Among the soil factors we find that the five with the highest average salience were soil pH (0.488), phosphorus ppm (0.487), potassium ppm (0.485), sulfate ppm (0.436), and percent organic matter (0.413) (Supplementary Figure 3 C).

Within the weather and management data, considering the average salience across the season (Supplementary Figure 3 D) five factors achieved an average salience greater than 0.140 – Total water (0.245), average solar radiation (0.198), maximum temperature (0.175), average wind direction (0.174), and estimated vapor pressure (0.173). The majority of factors had an average salience between 0.140 and 0.10 with six falling below this threshold – average soil temperature (0.095), maximum wind speed (0.084), average soil moisture (0.076), phosphorus applied (0.052) and potassium applied (0.033). Additionally, we find specific time points which appear to be salient broadly with the most salient region of time is within the first few days of planting, indeed 8 of the 10 days with the highest average salience are days 2-9 following planting.

### How is factor importance altered by inclusion of interactions?

The full CO model, in addition to performing best (albeit by a small margin), presents an opportunity to directly compare the influence of interactions between data modalities on the salience of factors because the single modality subnetworks are identical except for the prediction layer. The salience of genomic factors differs notably between the two networks (Supplementary Figure 3 B). Salience of PCs differs by as much as 0.432 (PC 24), with the difference in salience of the first 8 PCs (31% variance explained) ranging from 0.200 (PC1) to 0.309 (PC7). We find comparatively small differences in the salience of soil factors being between -0.011 and 0.0156 (Supplementary Figure 3 C).

In general, the salience map of the weather and management data features fewer broadly salient timepoints when interactions are included (Figure 3 A) than when they are not (Figure 3 B). The weather and management CO model contains a broadly salient time point around 25 days before planting and 6 days after planting. The SO model also appears to have peaks of salience around 150, 183, and 199 days after planting. When interactions are included the majority of the salient time points become less so with the exception of the peak 6 days after planting as highlighted through subtraction of the two salience maps (Figure 3 C).

## Discussion

### Assumptions, Potential Sources of Error, and Opportunities for Improvement

The results of this study are best understood with the data used and assumptions made kept in mind. The sole source of biological data in this study came from the Genomes to Fields Initiative (McFarland *et al*. 2020). The scale of this ambitious project increases the chances of data being absent or compromised due to equipment malfunction, logistical or procedural issues, and resource constraints. For example, many sites lack measurements for many soil properties across the seasons considered here, and the timing of fertilizer applications was absent in some cases. Our aim was to minimally filter the dataset while preventing missing or distorted values (many of which are not missing at random) from altering model accuracy and feature salience. We have aimed to reproducibly infer missing or aberrant values with relatively simple methods (e.g., imputation using linear models, KNN, etc.) but more sophisticated imputation techniques may have improved performance.

Alternatively, constraining the dataset to reduce the required imputation may have been an effective strategy. We elected to minimally filter observations because machine learning models, particularly deep learning, often benefit from having an abundance of data from which to learn feature relationships. For models where this is not the case, restriction of observations to the observations with the highest quality may be a preferable strategy. Note, however, that for distortions that are not randomly distributed, filtering may bias the sample and result in a model that appears to perform well but generalizes poorly (e.g., to sites similar to those with a preponderance of observations excluded).

Beyond including as many distinct locations and seasons as we could, we approximately balanced site by year groups through down sampling to avoid overfitting our DNNs to sites with more observations or biasing the selection of hyperparameters. This reduces the size of the dataset that can be used in training. Although outside the scope of this study, assessment of the sensitivity of DNNs to unbalanced group sizes, or exploration of alternate means of balancing groups (e.g., randomly *up sampling* small groups to equal the size of larger groups) would be valuable. Indeed, if balance were not a concern, or if it could be effectively achieved without discarding observations in some groups, one could potentially employ more strict data filtering without producing a dataset too small to benefit from machine learning.

Substantial effort was devoted to producing testing, training, and validation sets that would not lead to overconfidence in the accuracy of our models. To this end we kept observations within site-by-year groups in the same partition of the data. In effect, this prevents the model from being trained and tested on the same weather and management data.

Furthermore, except in cases where soil features are static from season to season, the model will not be trained and tested on observations with identical soil features. Proceeding in this manner rather than selecting observations at random for the testing set further reduces an already small number of weather and management conditions. Incorporating historical data (Washburn *et al*. 2021) or expanding the dataset to include data from other sources represent two possible avenues to incorporate a greater diversity of weather and management conditions without compromising the testing set.

Depending on the intended application of a model, one may be able to achieve higher performance through altering some of the above decisions or replacing random assignment with a targeted approach. For example, we assume that all group-by-year combinations are equally likely to be of interest. However, if we assume that the distribution of sites collected match those of interest for prediction (i.e., one is interested in predicting *any* future observation collected by G2F and the number of observations per field site are representative of future number of observations) then down sampling can be skipped, resulting a larger dataset. Similarly, with a narrower aim, e.g., prediction of yield within a specific region, testing or validation sets could be constrained to better select hyperparameters for or assess predictive accuracy of site-by-year combinations within that region.

In summary, our decision to include as much data as possible and to limit the possibility of overfitting to specific sites and seasons represent possible opportunities for improvement.

More sophisticated data imputation or more restrictive filtering, alternate means of balancing groups, and the incorporation of other data sources have the potential to improve model performance. Additionally, for more narrowly purposed models, non-random testing and training sets may represent a more accurate metric of predictive power, and indeed may deviate substantially from what we show here.

### Tradeoffs in Model Performance and Computational Resources

While the best performance was achieved with a deep neural network incorporating genomic, soil, weather, and management data, simple linear models with fixed effects often performed nearly as well (Figure 2). This is notable because tuning and training deep neural networks requires significant computational resources and time. For example, hyperparameter tuning in the machine learning models shown here took less than 24 hours to complete whereas tuning a single DNN sub-network took up several days. In the case of the best performing model this was repeated four times – once for each sub model.

**Figure 2.**
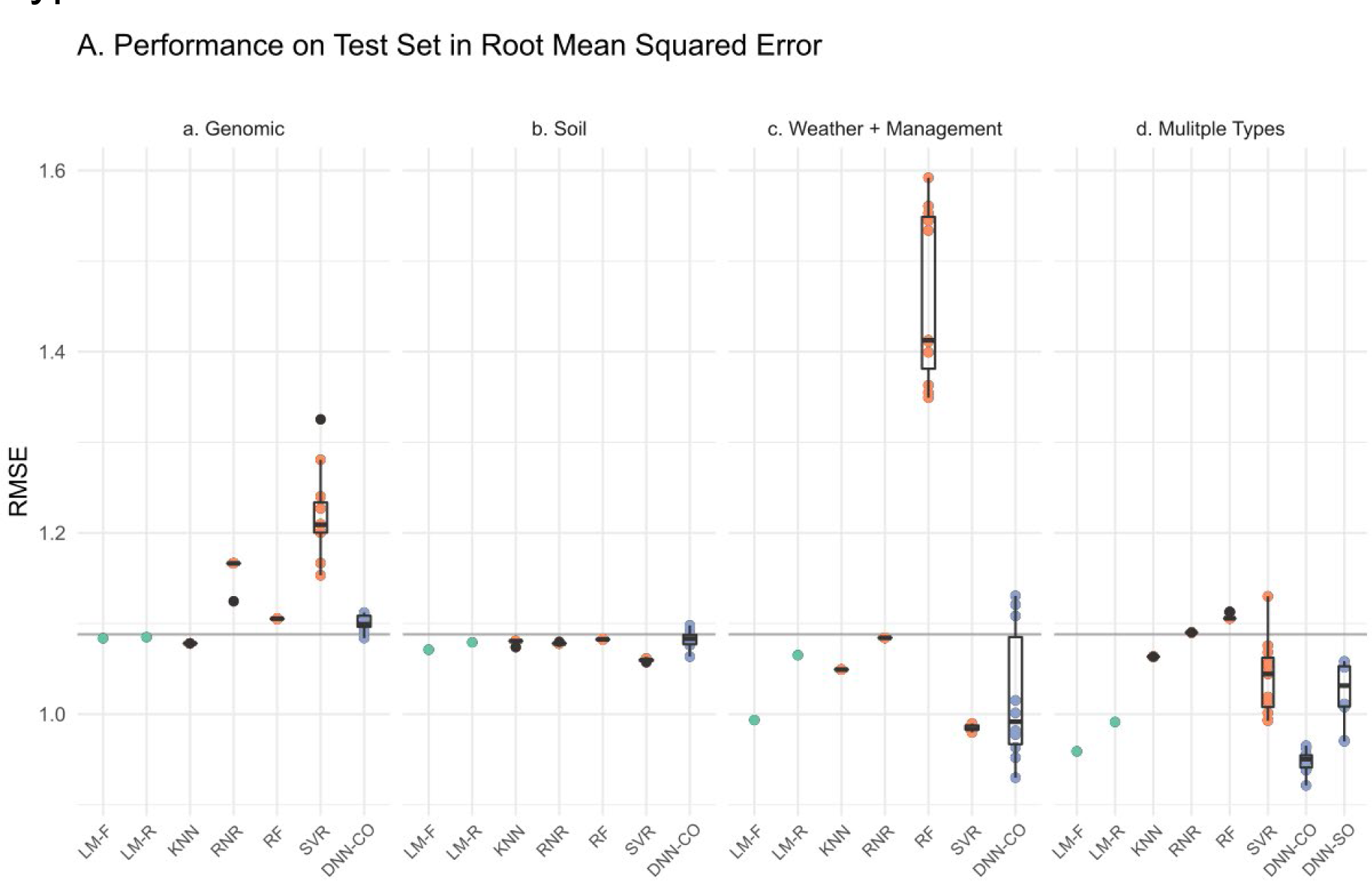
Model Performance Across Methodologies and Data Types **A.** The root mean squared error (RMSE) of the testing set is shown for each data grouping (panels a - d) and class of model. Lower values indicate better model performance. The horizontal gray line indicates the performance of an intercept model, i.e. using the mean of the training set yield as the prediction for all observations in the test set. For models that depend on a seed value the RMSE values for ten trials (evaluated on the same data) are shown and standard Tukey box plots are provided. In deep learning models random initialization of weights at the beginning of training result in different performance across trials. Three groups of models are shown, linear models (green), machine learning models (orange), and deep learning models (blue). Linear models are subdivided into those with exclusively fixed effects (LM-F) and those with random effects (LM-R). The best performing linear model is shown. For LM-F and LM-R respectively these are utilizing the first 8 PCs (explaining 31% of the variance) in **a**, utilizing all soil variables (for both LM-F and LM-R) in **b**, utilizing the five most salient weather/management factors (for both LM-F and LM-R) in **c**, and the first 8 genomic PCs, 5 most salient weather/management variables, and interactions between the two (for both LM-F and LM-R) **d**. Machine learning models used were K-Nearest Neighbors (KNN), Radius Neighbor Regression (RNR), Random Forest (RF), and Support Vector Regression with a linear kernel (SVR). Deep learning models are divided by whether they were part of the Consecutive optimization strategy (DNN-CO) or the Simultaneous optimization strategy (DNN-SO). Note that DNN-SO requires all data types and thus only appears in panel **d**.

By contrast, linear models, particularly those with only fixed effects, are quick to fit. They also outperformed many of the machine learning models, despite not undergoing extensive tuning for model structure. In cases where accuracy is not the sole factor under consideration, or where time or computational resources are limiting, simpler models may be “good enough” for the desired purpose.

### Usefulness of Consecutive Optimization in Hyperparameter Selection

We employed two strategies for hyperparameter optimization: consecutively optimizing (CO) hyperparameters for distinct “modules” of the network and simultaneously optimizing (SO) the network as a whole. CO reduces the range of possible combinations that are explored by allowing only one module to vary at a time. However, if two features in different data sets have a strong interaction effect (e.g., between genotype and weather patterns) then this approach will not necessarily allow for optimization to better capture this interaction. SO represents the reverse situation. With all features available, interactions between features in different tensors can be leveraged, but the hyperparameter space to explore is larger as all the hyperparameters are free to vary.

We find that the network resulting from CO substantially outperforms the one generated through SO. This should not be taken as a problem with SO *per se*. In other applications, or with a different optimization algorithm, it may prove to be a more efficient means of deriving a useful architecture. Furthermore, it is conceivable that SO is effective but that additional trials were required. The SO DNN architecture was selected based on 40 trials whereas the CO DNN architecture was selected based on 40 trials *for each module* (160 trials across the whole network) which confounds comparison. Selection of the training duration also warrants consideration. The SO model is capable of performing comparably to the CO model, but overfits more rapidly (Figure 1 B). Improved heuristics for selecting the training duration could increase usefulness of the SO model while reducing computational demands as well.

As a pragmatic matter, CO benefits from the capacity to tuning multiple modules at once. In our hands, total time spent tuning was driven more by modules with computationally intensive components (e.g. convolution layers) rather than the number of modules to optimize. This benefit is dependent on the tuning algorithm used. We used a bayesian optimization procedure which aims to produce useful hyperparameter combinations in fewer cycles than a simpler method such as grid approximation. However, because this method uses the performance of previously evaluated hyperparameters in selecting the next set, it does not permit parallelization in tuning a single network. If an optimization procedure that is conducive to parallelization were used (e.g., Hyperband or grid approximation) with enough computational resources this benefit would be non-existent.

Although we aimed to broaden the range of possible architectures relative to previous modeling on G2F data (Washburn *et al*. 2021), we constrained the overall structure to processing each tensor individually then allowing for interactions between the final layer of each module. Other options might include, for example, allowing an interaction module use both the first and final layers as input (instead of only the final one), or allow which layers were to be used to be tuned.

An additional option that we did not explore is aiming to inform the structure of the selected network based on known relationships between features. Similar to our decision to minimally transform and filter the data, we elected to avoid “nudging” the architecture of the network in any direction in order to allow the data to inform it instead. Informing the model architecture based on known relationships, analogous to incorporating a prior, remains an interesting and potentially fruitful avenue to pursue.

### Feature Importance

Similar to the results of previous modeling (Washburn *et al*. 2021), we find that no single data grouping provides sufficient information to disregard all others. We note that weather and management data does reduce error substantially relative to genetic and soil data, but the variation in performance is large (Figure 2). Only after integration of all data types do we see a relative reduction in error and consistency in this reduction.

Here we focus on salience in the weather and management data as it provided the best average performance when used without other datasets. We find that the total water applied to the field (including irrigation and rainfall, termed “WaterTotalInmm”) is the most influential factor for determining yield (Figure 3, Supplementary Figure 3 D). This is sensible from a biological standpoint and is in agreement with previous models. Previous DNNs developed with a subset of G2F data also identified precipitation as substantially influencing yield (Washburn *et al*. 2021). Linear modeling results find similar results and suggest a positive association between precipitation early in development and yield (Rogers *et al*. 2021). Additionally, in a recent study using a hybrid machine learning and crop growth model prediction system, the authors found that water related features (e.g. average drought stress, average water table in season) were important, although not as important as the trend in genetic and management improvements over time (Shahhosseini *et al*. 2021). The daily average of solar radiation (“SolarRadiationMean”) is the next most salient feature of this dataset, followed by the maximum temperature (“TempMax”) and the average wind direction (“WindDirectionMean”). A study employing a convolutional recurrent DNN to model county level data likewise found solar radiation and maximum temperature as important features and note an apparent increase in the importance of temperature near planting time (Khaki *et al*. 2020). A time dependent sensitivity can be observed in our model as well (Figure 3).

**Figure 3.**
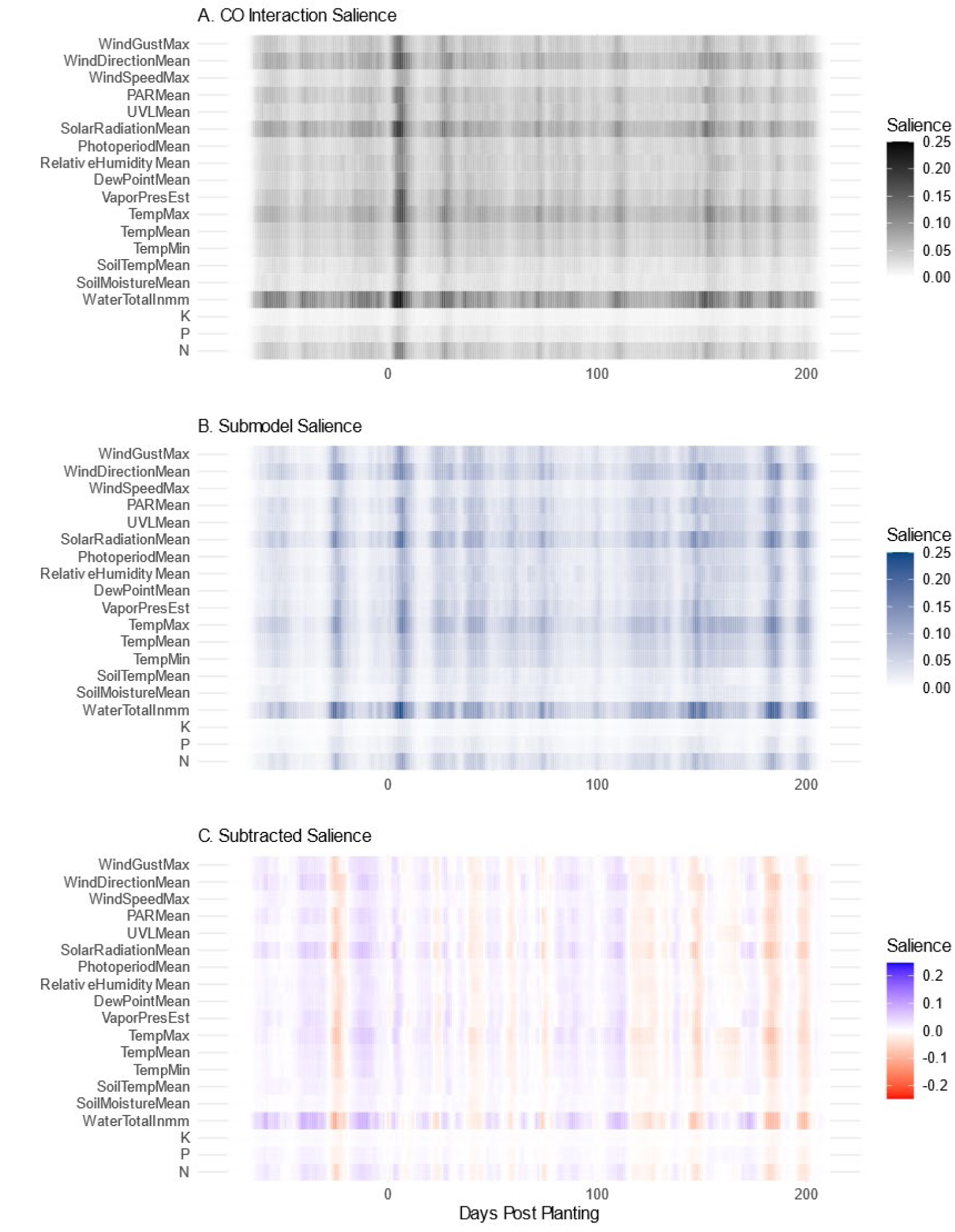
Influence of Interaction Effects on Feature Salience **A.** Average salience across all weather and management factors for each day considered. Interaction model values shown in black. **B.** The same values in **A.** are shown for the submodel in blue. Salience peaks shortly after planting in both models. The submodel contains additional peaks of salience prior to planting and near the end of the considered date range. **C.** Subtracted salience values for the interaction model and the submodel. The interaction-containing model appears to contain greater importance generally for certain features, e.g. irrigation and rainfall, represented as “WaterTotalInmm”. The difference between the two saliency maps indicates additional times of sensitivity in the submodel (approximately -25, +180, +195) that the interaction model is relatively insensitive to.

The relationship driving the high average salience of the average wind direction is not clear. This feature likely correlates with unrecorded variables. Assessment of the topology and geographical surroundings of each field site to suggest what this measure may be linked to lies outside the scope of this study.

With respect to management interventions, although addition of N, P, or K are not among the most salient weather and management features, we observe that nitrogen does have a mean salience comparable to relative humidity and photoperiod, while phosphorus and potassium are far lower. As noted in previously (Washburn *et al*. 2021) limited salience of fertilizers could be due to the quantities used being too low to exert a substantial effect, or alternatively application of these elements may be insufficiently variable to reveal the effect.

### Importance of GEM Interactions Accuracy in Feature Salience

Incorporating interactions between genetic, environmental, and management factors appears to have benefitted the accuracy of the resultant models. The CO DNN with interactions performed best (nRMSE 14.554%, RMSE 0.948) with the next best models, CO weather model and SO Models performing comparably (nRMSE 15.715%, RMSE 1.024). However, the two DNNs with interactions have far lower dispersion in RMSE, with standard deviations of 0.013 in the CO model and 0.035 in the SO model as compared with 0.074 in the CO weather model.

Interactions not only improve accuracy and model consistency across replicates, there appear to be changes in the salience of individual features as well. This is most apparent in considering weather and management features’ salience (Figure 3 C). Relative to the sub model, incorporating interactions appears to increase the salience of irrigation, although it is highly salient in both models (relative to other time series factors). Additionally, several broadly salient points in time, two of which are at the extreme end of the season, have diminished salience with the incorporation of interactions. This reduction is not uniform across all highly salient time points. A strong peak in salience shortly after planting is seen in both saliency maps which agrees with previously reported results (Washburn *et al*. 2021).

### Conclusions and Future Directions

The consecutively optimized deep neural network model developed here shows promise for complementing existing models for crop selection and improvement, as it produces more accurate estimates of yield than the other considered models. Of particular interest here is the capacity of this and other convolutional neural networks to incorporate change in environmental variables over time. This enables the generation of counterfactuals to examine the expected effect of different planting times (shifting the planting date of a site relative to the true value), planting in different sites, or planting under future possible climate scenarios. Additionally, the ability to generate such estimates would enable breeders to consider not only the expected yield of an individual cultivar but the expected *consistency* of yield as well.

For such a strategy to be adopted in genomic selection, further efforts are needed to validate the predictions such a model produces. This will necessitate incorporating of and validation on future data from the Genomes to Field Initiative (McFarland *et al*. 2020) or other large-scale experiments. The Genomes to Field Initiative and other organizations sponsor prediction competitions and other activities designed to advance this area of study.

Furthermore, applying the same model or the approach used to develop it to other crops would be a valuable step towards assessing its’ broad scale usefulness. This would also potentially implicate groups of crops for which the same model may be used through transfer learning, along with groups that require crop-specific models to be developed.

Additional improvements to accuracy that have the potential to transfer to modeling efforts for other crops include improved heuristics for epoch selection and training set construction. The simultaneously optimized model achieves a minimum error *lower* than our selected model (see Figure 2) and does so in far fewer epochs, but overfits much faster as well. If overfitting were preventable through a better heuristic for epoch selection than the one we employed, simultaneous optimization would have produced a better performing model that was simpler to generate. Training set construction is another opportunity for improvement with transferable utility. Here we took an aggressive approach ensuring approximately balanced groups, down sampling all groups with observations in excess of the smallest group in the test set. Deep neural networks tend to perform better with an abundance of data, so alternate approaches that retain more observations are of interest. In cases where there are few observations or model development is heavily constrained by computational resources or model development time, other models, especially linear regression models, may result in a model that performs nearly as well as a deep neural network.

Deep learning models do not result in parameters which are as readily interpretable as those of more standard statistical procedures and do not incorporate the physiology of the plant as mechanistic crop growth models do. These represent ongoing challenges and limit the scenarios in which a deep neural network may be useful. This can be partially addressed through how the data is represented (e.g. using non-PC transformed data), which has been explored for identification of genetic loci (Liu *et al*. 2019). Additionally, efforts to incorporate known relationships into a deep learning model’s structure have the potential to benefit accuracy and interpretability. Improvements in the capacity to represent genetic or physiological principles could allow for these methods to apply to a wider range of uses and address a broader set of questions.

## Author Contributions

Genomes to Fields experiments were coordinated and designed by Natalia DeLeon, David Ertl, Judith Kolkman, Dayane Cristina Lima, Danilo Moreta, James Schnable, and Maninder Singh. Field experiments were conducted, and data was collected and curated by Barış Alaca, Tim Bessinger, Natalia DeLeon, David Ertl, Sherry Flint-Garcia, Candice Hirsch, Joseph Knoll, Judith Kolkman, Dayane Cristina Lima, Danilo Moreta, James Schnable, Maninder Singh, Jason Wallace, Jacob Washburn, and Tecle Weldekidan. Joseph Gage provided novel genomic data. David Ertl and Maninder Singh contributed funding to the Genomes to Fields Initiative.

The computational study was designed by Daniel Kick and Jacob Washburn. Additional data cleaning and imputation was done by Daniel Kick, who developed the models and generated the figures. The manuscript was written by Daniel Kick and edited by Jacob Washburn, Jason Wallace, James Schnable, Judith Kolkman, and Daniel Kick.

## Acknowledgements

This research used resources provided by the SCINet project of the USDA Agricultural Research Service, ARS project number 0500-00093-001-00-D.

In addition to the contributions listed for the authors we would like to acknowledge those presently and historically involved in the Genomes to Fields Initiative, especially the following: Tim Bessinger, Martin Bohn, Edward Buckler, Natalia DeLeon, Jode Edwards, Sherry Flint-Garcia, Candice Hirsch, James Holland, Beth Hood, David Hooker, Shawn Kaeppler, Joseph Knoll, Sanzchen Liu, John McKay, Richard Minyo, Seth Murray, Rebecca Nelson, James Schnable, Rajan Sekhon, Maninder Singh, Peter Thomison, Addie Thompson, Mitch Tuinstra, Jason Wallace, Randy Wisser, and Wenwei Xu, who coordinated data collection during 2018 and 2019. Joseph Gage and Cinta Romay, who produced genotypic data. Alejandro Castro Aviles, Jode Edwards, David Ertl, Joseph Gage, James Holland, Dayane Cristina Lima, Bridget A McFarland, Christina Poudyal, Anna Rogers, Cinta Romay, Luis Samayoa, Kevin Silverstein, Tyson Swetnam, and Jacob Washburn, who curated the 2018 data. Ryan Timothy Alpers, Alejandro Castro Aviles, James Holland, Dayane Cristina Lima, and Bridget A. McFarland, who curated the 2019 data. Tecle Weldekidan made additional contributions to the project. Natalia de Leon, Dayane Lima, and Cinta Romay worked with Joseph Gage in production of genomic data.

## References

1. Anaconda Software Distribution, 2021 Anaconda Documentation.

2. Bache, S. M., and H. Wickham, 2020 magrittr: A Forward-Pipe Operator for R.

3. Bates, D., M. Mächler, B. Bolker, and S. Walker, 2015 Fitting Linear Mixed-Effects Models Using lme4. Journal of Statistical Software 67: 1–48.

4. Bergstra, J., D. Yamins, and D. Cox, 2013 Making a Science of Model Search: Hyperparameter Optimization in Hundreds of Dimensions for Vision Architectures, pp. 115–123 in Proceedings of the 30th International Conference on Machine Learning, edited by S. Dasgupta and D. McAllester. Proceedings of Machine Learning Research, PMLR, Atlanta, Georgia, USA.

5. Bradbury, P. J., Z. Zhang, D. E. Kroon, T. M. Casstevens, Y. Ramdoss et al., 2007 TASSEL: software for association mapping of complex traits in diverse samples. Bioinformatics 23: 2633–2635.

6. Buitinck, L., G. Louppe, M. Blondel, F. Pedregosa, A. Mueller et al., 2013 API design for machine learning software: experiences from the scikit-learn project, pp. 108–122 in ECML PKDD Workshop: Languages for Data Mining and Machine Learning,.

7. Chollet, F. and others, 2015 Keras. Couture-Beil, A., 2018 rjson: JSON for R.

8. Da Costa-Luis, C., S. K. Larroque, K. Altendorf, H. Mary, Richardsheridan, et al., 2022 tqdm: A fast, Extensible Progress Bar for Python and CLI. Zenodo. fuzzywuzzy, 2017 SeatGeek.

9. Harris, C. R., K. J. Millman, S. J. van der Walt, R. Gommers, P. Virtanen et al., 2020 Array programming with NumPy. Nature 585: 357–362.

10. Hornik, K., M. Stinchcombe, and H. White, 1989 Multilayer feedforward networks are universal approximators. Neural Networks 2: 359–366.

11. Hunter, J. D., 2007 Matplotlib: A 2D graphics environment. Computing in Science & Engineering 9: 90–95.

12. Inc, P. T., 2015 Collaborative data science.

13. Izrailev, S., 2021 tictoc: Functions for Timing R Scripts, as Well as Implementations of Stack and List Structures.

14. J. Liu, C. E. Goering, and L. Tian, 2001 A NEURAL NETWORK FOR SETTING TARGET CORN YIELDS. Transactions of the ASAE 44:.

15. Jarquin, D., N. de Leon, C. Romay, M. Bohn, E. S. Buckler et al., 2021 Utility of Climatic Information via Combining Ability Models to Improve Genomic Prediction for Yield Within the Genomes to Fields Maize Project. Front. Genet. 11: 592769.

16. Khaki, S., L. Wang, and S. V. Archontoulis, 2020 A CNN-RNN Framework for Crop Yield Prediction. Front. Plant Sci. 10: 1750.

17. Kibirige, H., G. Lamp, J. Katins, Gdowding, Austin et al., 2021 has2k1/plotnine: v0.8.0. Zenodo. Kubota, Y., 2021 tf-keras-vis.

18. Kurtzer, G. M., V. Sochat, and M. W. Bauer, 2017 Singularity: Scientific containers for mobility of compute (A. Gursoy, Ed.). PLoS ONE 12: e0177459.

19. Li, X., T. Guo, J. Wang, W. A. Bekele, S. Sukumaran et al., 2021 An integrated framework reinstating the environmental dimension for GWAS and genomic selection in crops. Molecular Plant S167420522100085X.

20. Liu, Y., D. Wang, F. He, J. Wang, T. Joshi et al., 2019 Phenotype Prediction and Genome-Wide Association Study Using Deep Convolutional Neural Network of Soybean. Front. Genet. 10: 1091.

21. Martín Abadi, Ashish Agarwal, Paul Barham, Eugene Brevdo, Zhifeng Chen et al., 2015 TensorFlow: Large-Scale Machine Learning on Heterogeneous Systems.

22. McFarland, B. A., N. AlKhalifah, M. Bohn, J. Bubert, E. S. Buckler et al., 2020 Maize genomes to fields (G2F): 2014–2017 field seasons: genotype, phenotype, climatic, soil, and inbred ear image datasets. BMC Res Notes 13: 71.

23. Messina, C. D., F. Technow, T. Tang, R. Totir, C. Gho et al., 2018 Leveraging biological insight and environmental variation to improve phenotypic prediction: Integrating crop growth models (CGM) with whole genome prediction (WGP). European Journal of Agronomy 100: 151–162.

24. Müller, K., 2020 here: A Simpler Way to Find Your Files.

25. O’Malley, T., E. Bursztein, J. Long, F. Chollet, H. Jin et al., 2019 KerasTuner. Pedersen, T. L., 2020 patchwork: The Composer of Plots.

26. Pedregosa, F., G. Varoquaux, A. Gramfort, V. Michel, B. Thirion et al., 2011 Scikit-learn: Machine Learning in Python. Journal of Machine Learning Research 12: 2825–2830.

27. R Core Team, 2021 R: A Language and Environment for Statistical Computing. R Foundation for Statistical Computing, Vienna, Austria.

28. Richardson, N., I. Cook, N. Crane, J. Keane, R. François et al., 2021 arrow: Integration to “Apache” “Arrow.”

29. Rogers, A. R., J. C. Dunne, C. Romay, M. Bohn, E. S. Buckler et al., 2021 The importance of dominance and genotype-by-environment interactions on grain yield variation in a large-scale public cooperative maize experiment (E. Akhunov, Ed.). G3 Genes|Genomes|Genetics 11: jkaa050.

30. Rogers, A. R., and J. B. Holland, 2021 Environment-specific genomic prediction ability in maize using environmental covariates depends on environmental similarity to training data (A. Lipka, Ed.). G3 Genes|Genomes|Genetics jkab440.

31. Samek, W., T. Wiegand, and K.-R. Müller, 2017 Explainable Artificial Intelligence: Understanding, Visualizing and Interpreting Deep Learning Models.

32. Seabold, S., and J. Perktold, 2010 statsmodels: Econometric and statistical modeling with python, in *9th Python in Science Conference*,.

33. Shahhosseini, M., G. Hu, I. Huber, and S. V. Archontoulis, 2021 Coupling machine learning and crop modeling improves crop yield prediction in the US Corn Belt. Sci Rep 11: 1606.

34. Simonyan, K., A. Vedaldi, and A. Zisserman, 2014 Deep Inside Convolutional Networks: Visualising Image Classification Models and Saliency Maps.

35. SingularityCE Developers, 2021 *SingularityCE 3.8.3*. Zenodo.

36. Tavenard, R., J. Faouzi, G. Vandewiele, F. Divo, G. Androz et al., 2020 Tslearn, A Machine Learning Toolkit for Time Series Data. Journal of Machine Learning Research 21: 1–6.

37. team, T. pandas development, 2020 *pandas-dev/pandas: Pandas*. Zenodo.

38. Technow, F., C. D. Messina, L. R. Totir, and M. Cooper, 2015 Integrating Crop Growth Models with Whole Genome Prediction through Approximate Bayesian Computation (I. De Smet, Ed.). PLoS ONE 10: e0130855.

39. Techtonik, A., 2015 wget 3.2.

40. Thornton, M. M., R. Shrestha, Y. Wei, P. E. Thornton, S. Kao et al., 2020 Daymet: Daily Surface Weather Data on a 1-km Grid for North America, Version 4.

41. Van Rossum, G., and F. L. Drake, 2009 *Python 3 Reference Manual*. CreateSpace, Scotts Valley, CA.

42. Virtanen, P., R. Gommers, T. E. Oliphant, M. Haberland, T. Reddy et al., 2020 SciPy 1.0: Fundamental Algorithms for Scientific Computing in Python. Nature Methods 17: 261– 272.

43. Washburn, J. D., E. Cimen, G. Ramstein, T. Reeves, P. O’Briant et al., 2021 Predicting phenotypes from genetic, environment, management, and historical data using CNNs. Theor Appl Genet 134: 3997–4011.

44. Waskom, M. L., 2021 seaborn: statistical data visualization. Journal of Open Source Software 6: 3021.

45. Westhues, C. C., G. S. Mahone, S. da Silva, P. Thorwarth, M. Schmidt et al., 2021 Prediction of Maize Phenotypic Traits With Genomic and Environmental Predictors Using Gradient Boosting Frameworks. Front. Plant Sci. 12: 699589.

46. Wickham, H., M. Averick, J. Bryan, W. Chang, L. D. McGowan et al., 2019 Welcome to the tidyverse. Journal of Open Source Software 4: 1686.

47. Zhou, D.-X., 2020 Universality of deep convolutional neural networks. Applied and Computational Harmonic Analysis 48: 787–794.

